# Superresolution Imaging Reveals the Spatial Organization of CD81 Microdomains in Regulating Membrane Signaling on Jurkat T Cell Microvilli

**DOI:** 10.1101/2024.12.07.627345

**Authors:** Neal T. Ramseier, Haoran Jing, Jesse Anderson, Ying S. Hu

## Abstract

Tetraspanin proteins are closely associated with high-curvature membrane structures and play key roles in organizing membrane domains and regulating membrane signaling in immune cells. However, their specific roles in regulating T cell membrane signaling, particularly within the microvilli often characteristic of these cells, remain poorly understood. Here, we used Jurkat T cells as a model system and investigated CD81 as a member of the tetraspanin family. Using total internal reflection fluorescence (TIRF) microscopy and structured illumination microscopy (SIM), we identified an enrichment of the tetraspanin CD81 microdomains along the actin-rich membrane microvilli. At the distal end of the microvilli, SIM images revealed the spatial colocalization of CD81 with T cell receptors (TCR) and CD63, implying a potential role for CD81 in regulating TCR signaling in conjunction with CD63. Spatial analysis of CD81 and CD63 microdomains from the dual-color SIM data revealed their preference for associating with each other. Cluster analysis of direct stochastic optical reconstruction microscopy (*d*STORM) data revealed that *in vitro* T cell activation results in reduced domain sizes and increased domain separation of CD81. These findings provide visual evidence of the spatial organization and rearrangement of CD81 on the T cell microvilli, highlighting its potential role in signal regulation on specialized membrane protrusions.

## Introduction

Tetraspanins are membrane proteins that associate with each other to form tetraspanin-enriched microdomains (TEMs) on the plasma membrane, playing key roles in the organization of the membrane architecture (*1–3*). CD81 represents a tetraspanin protein consisting of four transmembrane domains and is localized in TEM on the cell surface. CD81 mediates numerous cellular functions, including cellular adhesion, intracellular signaling, proliferation, and regulation of T cell morphology (*4, 5*). In addition, CD81 has been reported to localize on the microvilli in endothelial cells (*6*) and is known for regulating membrane protrusions (*7*). CD81 has been demonstrated to localize on the microvilli of B cells and monocytes and play a role in microvilli regulation in primary mouse peripheral blood mononuclear cells (*7*).

In T cells, membrane microvilli are dynamic, actin-associated structures that facilitate environmental probing and sensing (*8–10*). The enrichment of signaling proteins allows microvilli to survey antigens on antigen-presenting cells and form complexes for subsequent signaling processes (*11–13*). The signaling function of T cell microvilli is suggested by the clustering of T cell receptors (*12–14*), as well as the release of T cell microvilli particles (TMP) containing adhesion molecules and cytokines (*11, 15, 16*). Given the known association of tetraspanins with high-curvature membrane structures, understanding the nanoscale spatial organizations of tetraspanin proteins on the microvilli enhances our understanding of the signaling regulation mechanism at these specialized membrane structures.

Notably, electron microscopy (EM) using immunogold staining has previously revealed the presence of CD81 on Jurkat T cell microvilli (*17*). CD81 has been reported to regulate T cell activation (*18–20*) and is known to associate with CD4 and CD8 (*21, 22*). Moreover, CD81 has been shown to act as a costimulatory molecule to activate T cells when stimulated with CD3 (*23– 25*). While these studies point to the potential role of signal regulation, the nanoscale distribution and spatial arrangements of CD81 concerning other membrane proteins on the microvilli have not been visualized. Such information enhances the understanding of T cell signaling in specialized membrane protrusions.

In this study, we employed fluorescence and superresolution imaging to examine the spatial distribution of the CD81. We observed a CD81 enrichment with TEMs on Jurkat T cell microvilli. We investigated the spatial relationship of CD81 with T cell receptors (TCR) and CD63 along and on the distal end of microvilli. We further revealed the ultrastructure of CD81 TEMs and their spatial redistribution upon *in vitro* activation.

## Results

### CD81 enrichment on T cell microvilli revealed by fluorescence and superresolution imaging

F-actin has been shown to be involved in the formation of cytoskeletal protrusions in T cells (*11, 12*), dendritic cells (*26*), and other cell types (*27, 28*). To assess the localization of CD81 on the microvilli, we immobilized and fixed Jurkat T cells, then co-stained CD81 and F-actin (**Figure 1A**). **Figure 1B** displays total internal reflection fluorescence (TIRF) images of CD81 and F-actin colocalizing on microvilli structures protruding from Jurkat T cells. The F-actin distribution shows a preference toward the periphery of the cell, in conjunction with CD81. The peripheral F-actin distribution is consistent with previous reports of F-actin staining on T cells (*9*). The magnified views of the microvilli suggest a punctated distribution of CD81 compared to the continuous appearance of F-actin, revealing the presence of CD81 TEMs on the microvilli. CD81 is localized both on and next to the actin fibers.

**Figure 1:**
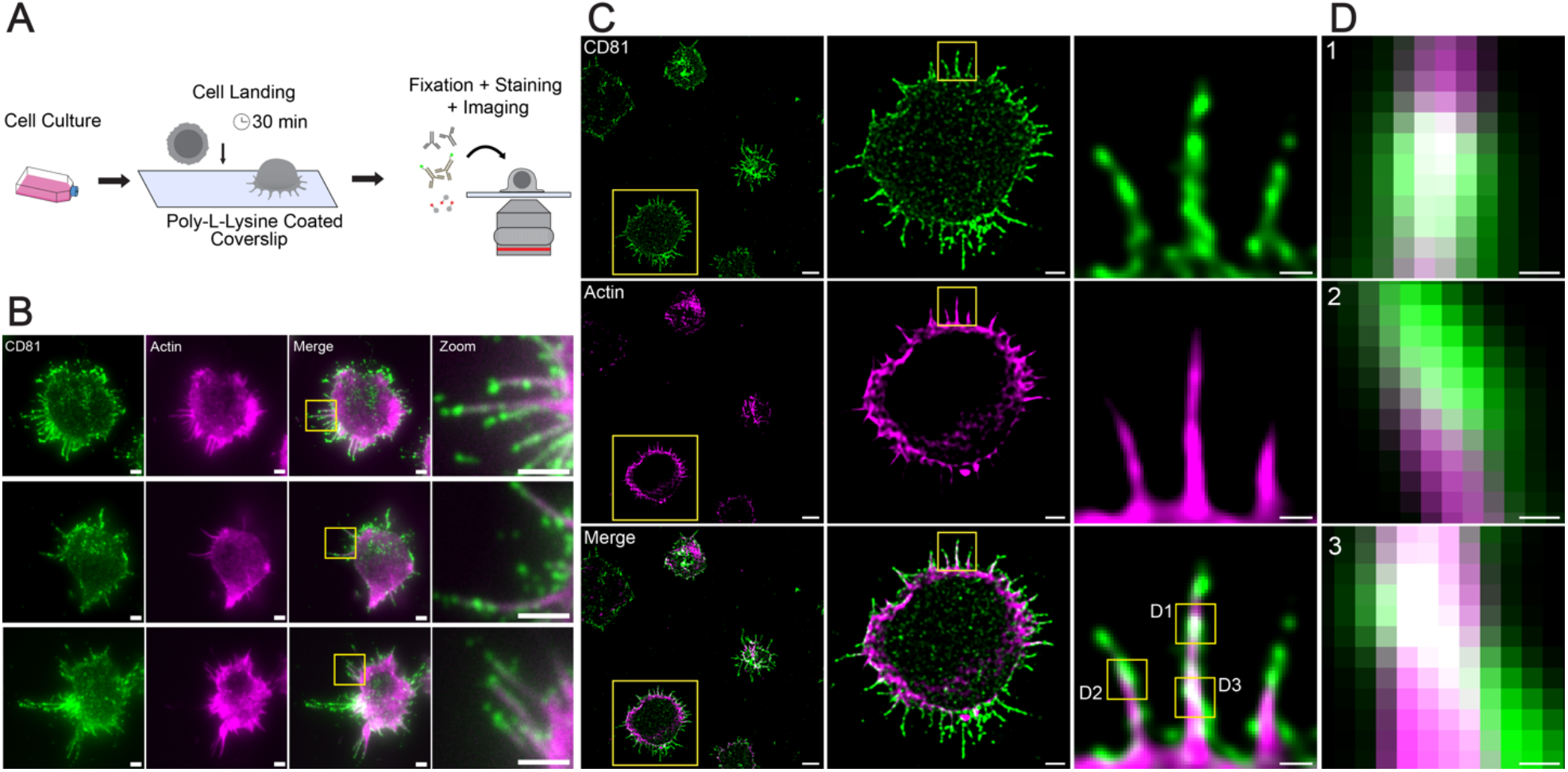
Fluorescence superresolution microscopy demonstrated CD81 localization on Jurkat T cell microvilli. (**A**) Schematic depicting the immobilization, fixation, and staining of Jurkat T cells for imaging. (**B**) TIRF immunofluorescence images of CD81 (green), F-actin (magenta), merged images of CD81 and F-actin, and magnified views of the box across Jurkat T cells. (**C**) DeepSIM images of CD81 (green), F-actin (magenta), and merged images of CD81 and F-actin across Jurkat T cells. Magnified views are shown from left to right. (**D**) Magnified views of the boxed regions in panel **C**. Scale bars: 5μm (**C left**), 2μm (**B, C middle**), 500 nm (**C right**), and 100 nm (**D**).

To obtain higher spatial resolution, we performed structured illumination microscopy using DeepSIM (CrestOptics) and confirmed the distribution of CD81 clusters along the microvilli (**Figure 1C**). In contrast, the intracellular distribution of CD81 was much lower. Colocalization analysis using the intensity correlation quotient (ICQ) (*29*) revealed an ICQ value of 0.190 ± 0.017 (mean ± SEM, *n* = 15) for the microvilli (**Figure S1**), suggesting a positive correlation between CD81 and F-actin. High-magnification views in **Figure 1C** further revealed the segmented TEMs of CD81, which were identified on the microvilli of most of the Jurkat T cells. Furthermore, the higher resolution revealed spatial arrangements of CD81 in the context of F-actin (**Figure 1 D**). We employed volumetric superresolution imaging and confirmed the prevalence of CD81-enriched microvilli across the entire Jurkat T cell surface **(Figure S2**).

Importantly, we observed CD81 TEMs on the microvilli with or without membrane permeabilization. Previous reports have shown tetraspanins to remain associated under non-stringent detergent conditions (*30, 31*). A comparison between fixatives (4% paraformaldehyde (PFA) or 4% PFA + 0.1% glutaraldehyde (GA)) with and without permeabilization using saponin all revealed similar phenotypes of CD81 intensity variation along the microvilli (**Figure S3**). The observation of this phenotype in all four conditions suggests the segmented distribution of CD81 is not due to permeabilization agents washing away the membrane enclosing the microvilli. The heterogeneous distribution of CD81 signal intensity suggests the spatial clustering of CD81 molecules into TEMs along the microvilli.

### TCRs are spatially associated with CD81 TEMs on the microvilli

CD81 is known to regulate TCR signaling (*20*). Considering the observed enrichment of CD81 at the microvilli and previously reported accumulation of TCRs on and at the distal ends of microvilli (*11, 13, 14, 32*), we investigated the spatial relationship between these two proteins. The dual-color SIM image of Jurkat T cells immunostained for CD81 (green) and TCRs (blue) in **Figure 2A** revealed a strong correlation between the spatial distribution of these two proteins. Notably, TCRs present as punctated domains similar to the CD81 TEMs. The density of TCR microdomains appears to be sparser than CD81 TEMs.

**Figure 2:**
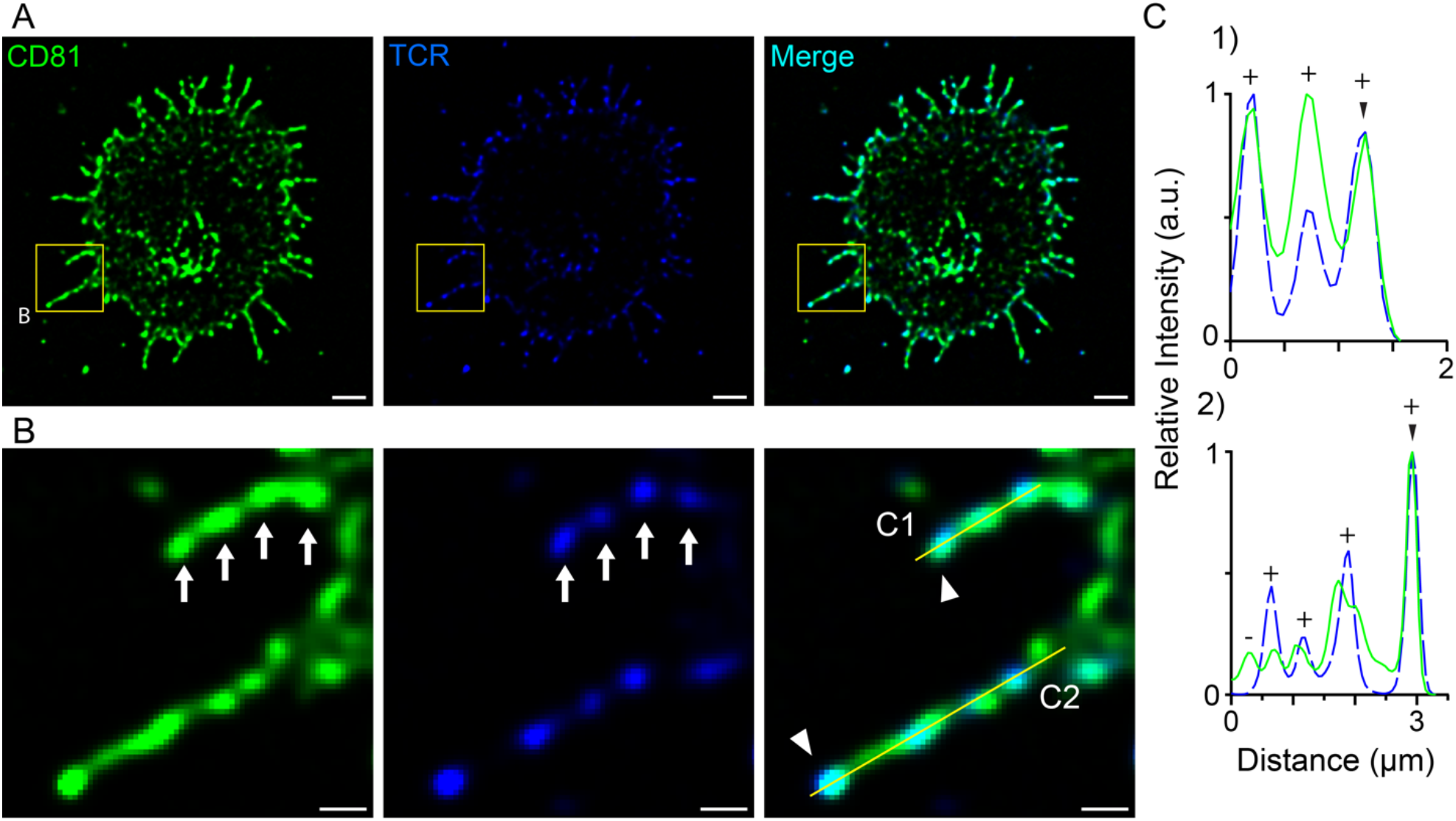
TCR colocalizes with CD81 on the microvilli. (**A**) DeepSIM images of CD81 (green), TCR (blue), and merged image of CD81 and TCRs across a Jurkat T cell. (**B**) Magnified view of boxed region in panel **A**. Arrows point to CD81 TEMs that contain TCR. Triangles in the right panel point to the distal end of the microvilli. (**C**) Line profiles displaying the relative intensity of CD81 (solid green) and TCR (dashed blue). (+) denotes TCR colocalization with CD81, and (–) denotes CD81 without TCR. Triangles point to the distal end of the microvilli. Scale bars: 2 μm (**A**) and 500 nm (**B**).

The magnified views displayed strong colocalization (cyan) of TCRs with CD81 along the microvilli (**Figure 2B)**. The white arrows in **Figure 2B** point to similar morphology and spatial distributions of CD81 and TCR microdomains. The line profiles in **Figure 2C** further confirm the spatial correlation between the two. The TCR puncta associated with CD81 puncta are denoted as + on the line profiles. Notably, not all CD81 TEMs contain TCRs, as denoted by – on the line profile. TCR and CD81 colocalize at the distal end of the microvilli, as marked by triangles. These results demonstrate a close spatial relationship between TCRs and CD81 TEMs, suggesting that CD81’s proximity to TCRs facilitates its regulatory role in T cell signaling across the microvilli surface.

### CD63 TEMs are less dense than CD81 TEMs and exhibit strong colocalization with TCRs at the distal ends of microvilli

We next investigated CD63 in the context of CD81 on the T cell microvilli, as tetraspanins are known to associate with one another to form TEMs (*33*). CD63 has been observed to be present on micro-protrusions in activated platelets (*34*). Similar to CD81, CD63 has been reported to be involved in T cell activation and can be used as a costimulatory target for activation with CD3 (*23, 35*). SIM images of co-stained CD81 and CD63 revealed that CD63 presents as punctated TEMs and is not as abundant as CD81 on the Jurkat T cells (**Figure 3A**). We observed a preference for CD63 association with CD81 toward the periphery of the cell, including the microvilli. Not all CD63 TEMs are spatially associated with CD81 TEMs, but they display strong association on several microvilli, as seen in the magnified views of **Figure 3A**. The line profiles in **Figure 3B** show the relative intensity of CD63 and CD81, with a CD63 peak at the distal end of the microvilli colocalizing with CD81, denoted by the triangles. The distal ends of the microvilli show spatial colocalization between CD81 and CD63 (**Figure 3 B1-3**). It is important to note that DeepSIM images of 0.2 µm Tetraspeck^TM^ microspheres on a coverglass in 488 nm, 561 nm, and 640 nm channels displayed complex chromatic shifts, which varied both among the microspheres and across the 488 nm, 561 nm, and 640 nm channels (**Figure S5**). We observed chromatic shifts as large as approximately 120 nm, which is convolved with the multicolor superresolution images obtained by DeepSIM.

**Figure 3:**
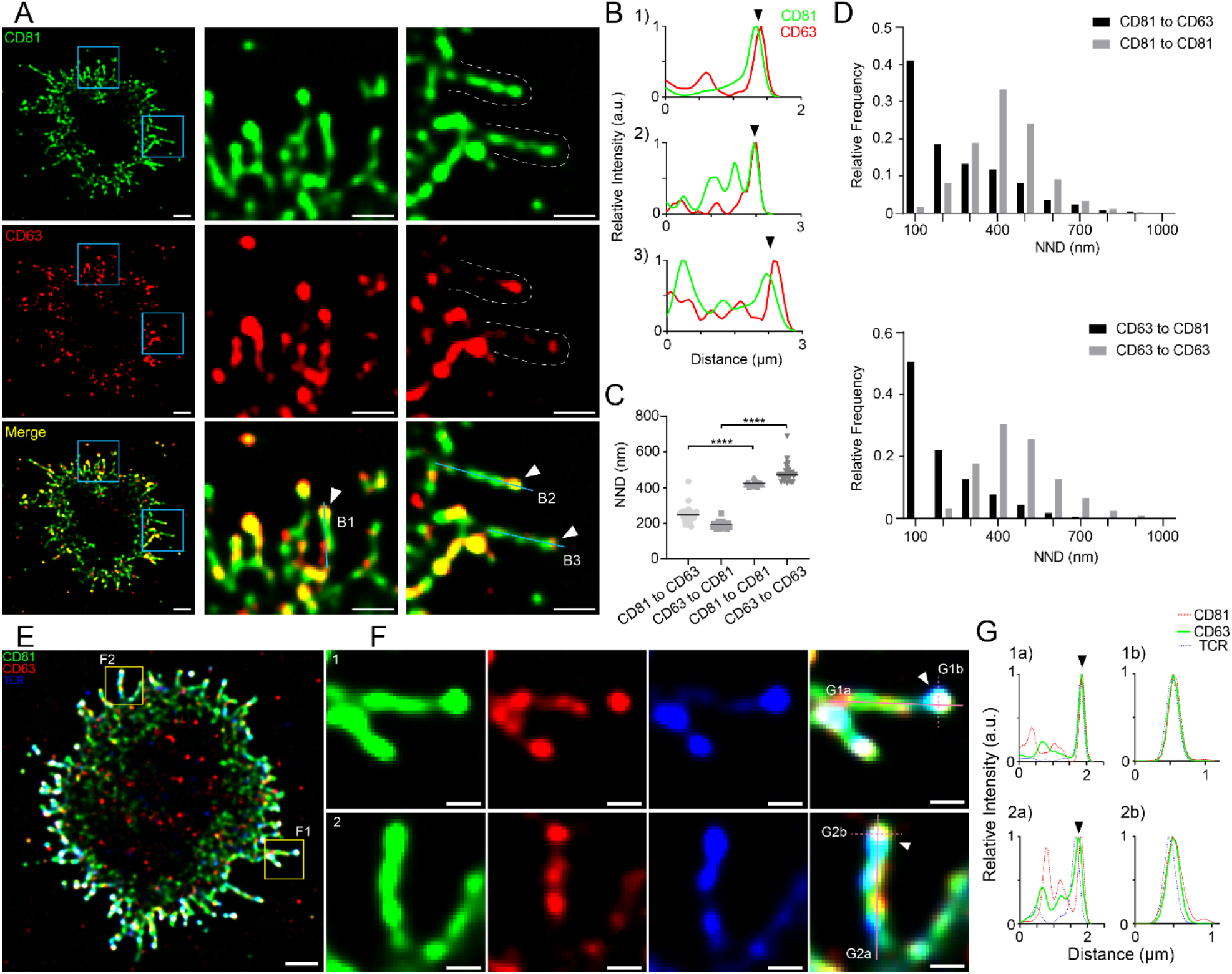
CD63 associates with CD81 and colocalizes with CD81 and TCR on distal ends of microvilli. (**A**) DeepSIM images of CD81 (green), CD63 (red), and merged images of CD81 and CD63 across a Jurkat T cell. The magnified views of the boxed region from the left panel are shown in the center and right panels. The two microvilli in the right panel are outlined with the dashed lines. (**B**) Line profiles for B1-3 labeled in panel **A**. Triangles denote the distal end of the microvilli. (**C**) Comparison of the per-cell mean nearest neighbor distance (NND) between the TEMs consisting of CD81 or CD63 along the microvilli and cell periphery (*n* = 28). *n* represents the number of cells, and each data point represents the mean NND from one cell. Bar displays the mean. Statistical significance was evaluated using a Student’s *t*-test (^****^ p < 0.0001). (**D**) The distribution of NNDs pooled from all cells characterized in C): CD81 to CD63 (*n* = 10,428), CD81 to CD81(*n* = 10,428), CD63 to CD81 (*n* = 8,355), and CD63 to CD63 (*n* = 8,355). *n* represents the number of NND measurements between two TEMs. (**E**) DeepSIM image of CD81 (green), CD63 (red), and TCR (blue) across a Jurkat T cell. (**F**) Magnified views of the boxed region in panel **E**. Triangles denote the distal end of the microvilli. (**G**) Line profiles of the longitudinal and transverse profiles marked in the right panel of **F**. Triangles denote the distal end of the microvilli. Scale bars: 2 μm (**A left, E**) and 500 nm (**A middle & right, F**).

We further quantitatively characterized the spatial relationship between CD81 and CD63. We analyzed the nearest neighbor distance (NND) between intensity maxima of CD63 and CD81, comparing CD63 to CD81, CD81 to CD63, CD81 to CD81, and CD63 to CD63 (**Figure S4**). Interestingly, we observed that the peaks of CD81 are closer to the peaks of CD63 than to other peaks of CD81, and the peaks of CD63 are similarly closer to CD81 peaks than to other peaks of CD63 (**Figure 3C**). The difference in distance between CD81 or CD63 to itself and the other was found to be statistically significant (*n* = 28, *p* < 0.0001) using a Student’s *t*-test. Specifically, both CD81 to CD63 and CD63 to CD81 display a decreasing relative frequency as the NND values increase, while CD81 to CD81 and CD63 to CD63 relative frequencies display a normal distribution, peaking at around the mean value (**Figure 3D**). This trend persists even after excluding the 100 nm NNDs between CD81 and CD63, due to uncertainties arising from the chromatic shifts mentioned earlier. The findings indicate a strong correlation between the organization of CD63 and CD81 on the microvilli and cell periphery. Colocalization analysis of the cells further supports this notion with a calculated ICQ value of 0.334 ± 0.008 (mean ± SEM, *n* = 28).

Given the colocalization of CD81 with CD63 at the distal ends of microvilli and the shown colocalization of CD81 with TCRs, we immunostained Jurkat T cells for CD81 (green), CD63 (red), and TCR (blue). The DeepSIM image in **Figure 3E** revealed strong colocalization (white) between all three proteins at the distal ends of the microvilli (**Figure 3F**). The magnified views on the right and the line profiles in **Figure 3G** further display this colocalization of all three proteins at the distal ends of the microvilli. These results suggest a close spatial relationship between CD81, CD63, and TCRs, providing visual evidence of CD81 and CD63’s cooperative roles in regulating TCR signaling on the microvilli surface through physical proximity.

### *d*STORM reveals nanoscale spatial rearrangements of CD81 TEMs across the microvilli after *in vitro* activation

We employed *d*STORM (*36, 37*) to examine the nanoscale ultrastructure of CD81 TEMs on Jurkat T cell microvilli. **Figure 4A** presents a *d*STORM image of CD81 on a control Jurkat T cell. The accumulation of CD81 localizations at the periphery of the cell, particularly along the microvilli, is shown in the magnified views in **Figure 4A**. We observed CD81 TEMs consistent with the DeepSIM data. The high resolution of *d*STORM enables the measurement of the microvilli width and was found to be 100-300 nm (**Figure 4B, C**). This observation is consistent with previous literature, factoring in the antibody size and localization precision (*38*).

**Figure 4:**
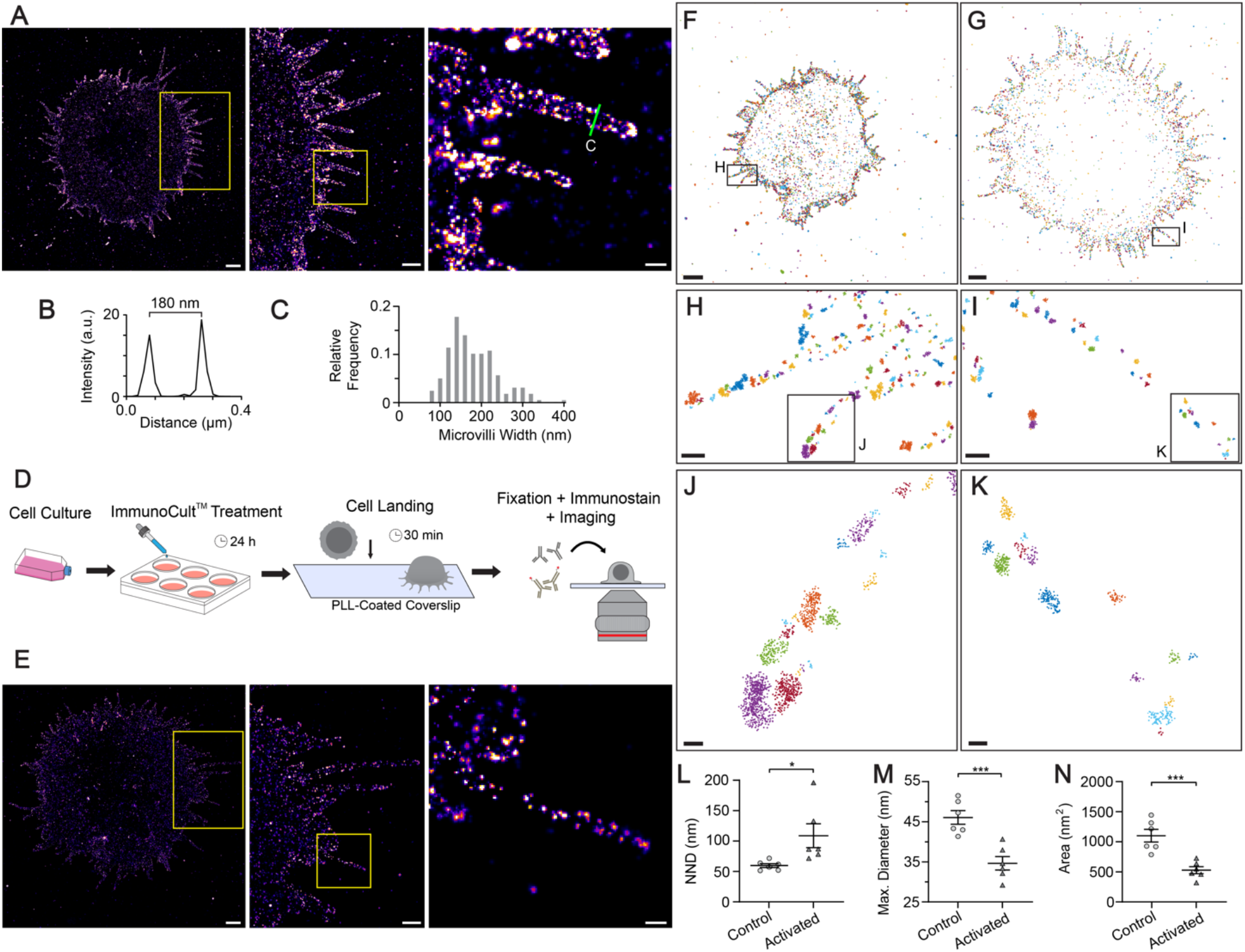
*d*STORM imaging reveals nanoscale clustering and rearrangement of CD81 TEMs on the microvilli with *in vitro* activation. (**A**) *d*STORM image of CD81 across a Jurkat T cell. Magnified views of the boxed region are shown to the right. (**B**) The line profile shown in panel **A (right)** depicting the width of a microvillus. (**C**) Distribution of microvilli widths. (**D**) Schematic depicting ImmunoCult^TM^ treatment, landing, fixation, and immunostaining of 24-h activated Jurkat T cells for imaging. (**E**) A *d*STORM image of CD81 across a 24-h activated Jurkat T cell. Magnified views of the boxed region are shown to the right. (**F**) Clustering analysis by DBSCAN on a control Jurkat T cell. (**G**) Clustering analysis by DBSCAN on a 24-h activated Jurkat T cell. (**H**) Magnified view of the box shown in panel **F**. (**I**) Magnified view of the box shown in panel **G**. (**J**) Magnified view of the box shown in panel **H**. (**K**) Magnified view of the box shown in panel **I**. (**L**) Comparison of the NND of CD81 clusters between control and 24-h activated cells. (**M**) Comparison of the maximum diameter of CD81 clusters between control and 24-h activated cells. (**N**) Comparison of the cluster area of CD81 clusters between control and 24-h activated cells. In **L-N**, each data point represents the mean value from the clusters in each cell. *n* = 6 for control cells and *n* = 6 for 24-h activated cells. Bar graphs display mean and SEM. ^***^*p*<0.001, ^*^*p*<0.05. Scale bars: 2 μm (**A(left), E(left), F, G**), 1 μm (**A(middle), E(middle)**), 250 nm (**A(right), E(right), H, I**), and 50 nm (**J, K**).

Given CD81’s role in T cell activation and signaling regulation, we assessed the extent to which *in vitro* activation influences the CD81 nanoscale ultrastructure. A previous study found that *in vitro* activation results in an immediate reduction in CD81 expression on the cell surface, with an over 50 % decrease at 24 hours using flow cytometry (*17*). To observe the maximal difference in CD81 surface expression due to activation, we incubated Jurkat T cells with ImmunoCult^TM^ for 24 hours. The treated cells were landed and immobilized on a PLL-coated surface, then fixed, immunostained for CD81, and imaged by *d*STORM using the same experimental and image reconstruction conditions (**Figure 4D)**. Compared to the control cells, 24-h ImmunoCult^TM^activated cells displayed sparser CD81 localization densities along the microvilli (**Figure 4E**). The magnified views of the microvilli in **Figure 4E** compared to **Figure 4A** further highlight this difference. The differential CD81 distribution was validated with TIRF and SIM images of CD81 on control and 24-h activated Jurkat T cells (**Figure S6**). We further confirmed the presence of microvilli on the activated Jurkat T cells using a membrane dye (**Figure S7**).

To quantitatively analyze the clustering, we employed a density-based spatial clustering of applications with noise (DBSCAN) (*39*) algorithm to identify individual clusters of CD81 (**Figures 4F,G**). The magnified views of the control cell (**Figure 4H,J**) and 24-h activated cell (**Figure 4I,K**) display a difference in cluster sparsity between the two conditions. To this end, we quantified the microvilli CD81 clusters’ NND. We included only clusters on the microvilli (**Figures S8, S9**) and calculated the mean cluster NND from each cell for comparison. In activated cells (*n* = 6), the mean NND significantly increased compared to control cell (*n* = 6, *p* = 0.0328) (**Figure 4L**). These results quantitatively confirm the increased sparsity of clusters after activation. The higher-magnification views in **Figures 4J, K** indicate smaller CD81 clusters after activation. To quantify the observed difference in cluster size, we calculated the maximum diameter (MD) of each cluster along the microvilli, defined as the maximum distance between two points within a cluster, as well as the total area of each cluster bounded by the outer points. The mean MD of the clusters from each cell in the control and 24-h activated showed a decrease after activation, found statistically significant (*p* = 0.0007) **(Figure 4M)**. The decrease in cluster size after activation is confirmed by an observed decrease in the cluster area, also found statistically significant (*p* = 0.0008) (**Figure 4N**). Taken together, the cluster analysis of the *d*STORM images of control and activated cells indicates a spatial rearrangement of CD81 TEMs following activation and provides further detail of the ultrastructure of CD81 TEMs on Jurkat T cell microvilli.

## Discussion

We observed an enrichment of CD81 and colocalization with F-actin at the microvilli, confirming its designation as a T cell microvilli biomarker. The punctated microdomains observed in the DeepSIM images correlate to the areas with larger clusters or multiple clusters in close proximity seen in the DBSCAN analysis. These areas of high CD81 fluorescence intensity and localization density could likely be attributed to the organization of CD81 into TEMs (*40, 41*), and tetraspanins have been shown to regulate spatiotemporal distributions of membrane proteins to regulate immune cell signaling (*42*). Of note is that TEMs are not unique to T cells and are found in different cell types (*43, 44*). CD81 in various cell types has been implicated in viral infection, including human immunodeficiency virus (*44*), Influenza (*45, 46*), and Hepatitis C (*47, 48*). Influenza viral budding occurs in membrane regions highly enriched with CD81, which may utilize a mechanism similar to the release of certain EVs, particularly microvesicles, from cells (*11, 17*).

CD81 has been shown to regulate the signaling regarding TCRs (*18, 19*). Previous studies reported CD81 internalization after activation (*17*) and the migration of CD81 to the immunological synapse (IS) during its formation (*18, 49*). The past findings, in addition to our observations of CD81 colocalizing with TCRs on the microvilli, strongly implicate CD81 as a regulator of T cell activation. Our observation of CD63 colocalized with CD81 at the distal ends of microvilli, in addition to its role as a costimulatory molecule (*35*), implicates CD63’s participation in the signaling regulation of T cell activation (**Figure 5A**). The exact role of CD63 in T cell activation signaling remains to be understood; however, its localization with CD81 and TCRs provides direct evidence of spatial association between these molecules. As CD63 can serve as a costimulatory molecule for T cell activation, the localization would likely be near TCRs. The presence of both CD81 and CD63 further suggests the formation of TEMs at the distal ends of microvilli where TCR is enriched. Future studies looking into other proteins within these domains would provide further insight into the exact organization of the microvilli around TCR.

**Figure 5:**
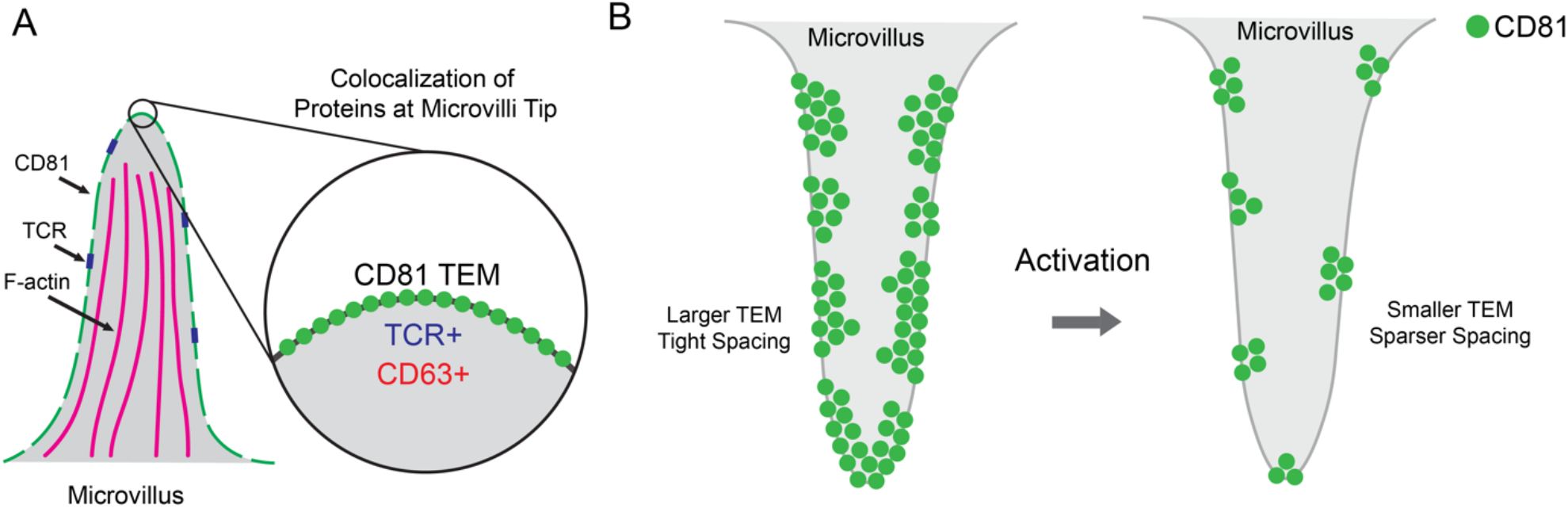
CD81 colocalization with CD63 and TCR on the distal end of actin-rich Jurkat T cell microvilli and CD81 TEMs rearrange after *in vitro* activation. (**A**) Illustration depicting TEMs of CD81 along the actin-rich microvilli. The magnified view of the distal end of the microvillus shows CD81 TEMs colocalize with CD63 and TCR. (**B**) Illustration depicting the spatial rearrangement of CD81 TEMs on the microvilli of Jurkat T cells after activation, highlighting a reduction in TEM size and an increase in TEM separation.

In the Jurkat T cells we imaged, the distribution of CD81 on the microvilli of CD3/CD28 treated cells appeared sparser than the microvilli of the control (non-manipulated) Jurkat T cells (**Figure 5B**). Quantitative analysis of CD81 clusters identified using DBSCAN revealed smaller clusters and a sparser distribution of clusters on the microvilli of 24-h activated cells compared to control cells. This observation is consistent with a previous report noting a rapid decrease in surface CD81 expression, measured with flow cytometry after treatment with CD3 and CD28 (17). The observed decrease in CD81 density on the Jurkat T cell microvilli may be linked to its role as a regulatory molecule regarding the TCR. CD81 has been shown to suppress T cell signaling, likely acting as a safety mechanism for accidental triggering (*19*) and as a co-stimulatory molecule to TCR stimulation (*24*). After 24-h of activation, this regulatory and stimulatory mechanism may no longer be needed as the needs of the cell change, as indicated by a reduction in CD81 mRNA levels after 9-h of activation (*17*). The spatial reorganization of CD81 on the T cell microvilli in response to T cell activation suggests that specialized membrane protrusions are involved in receptor-mediated signaling and potentially downstream signaling cascade. Specifically, the enrichment of CD81, its previously reported role in T cell activation, and the enrichment of TCR on the microvilli suggest the microvilli as the major signaling hub for T cell activation. The presence of CD63 on the microvilli and its known role as a costimulatory molecule further supports this concept.

Our study’s limitations include using Jurkat T cells as a model system and the cells’ attachment to the glass surface. The cellular attachment to the surface and the absence of a permeabilization agent may affect the accessibility of antibodies to the membrane and microvilli directly underneath the cell. Future studies employing live-cell imaging with fluorescent proteins fused to CD81 could provide further insights into the spatial distribution of CD81 in the context of the temporal behavior of T cell microvilli, as microvilli are dynamic structures. Further studies to determine the exact roles of CD81 and CD63 in T cell activation and signaling will provide more insight into the roles of tetraspanins in regulating the T cell plasma membrane.

In conclusion, we identified CD81 as a microvilli marker in Jurkat T cells through its enrichment on these membrane protrusions. CD81 TEMs colocalize with TCRs along the microvilli and with CD63 and TCRs at the distal end of microvilli, providing evidence of CD81’s involvement in regulating T cell activation with CD63. The *d*STORM clustering analysis provides a quantitative understanding of the spatial redistribution of CD81 following T cell activation. These results suggest CD81 TEMs regulate T cell signaling on the microvilli through spatial association.

## Materials and Methods

### Reagents

Fetal Bovine Serum (F0926-500ML), tris(hydroxymethyl)aminomethane (252859-500G), sodium chloride (S3014-1KG), D-(+)-Glucose (G8270-1KG0), glucose oxidase from *Aspergillus niger* (G7141-50KU), catalase from bovine liver (C40-100MG), 0.01% Poly-L-Lysine Solution (P4707-50ML) and 2-mercaptoethanol (M6250-100ML), saponin (S-4521) were purchased from MilliporeSigma. RPMI (11875093), penicillin-streptomycin (15140-122-100 mL), DPBS (14190-144-500 mL), Cellvis chambered cover glass (C8-1-N), goat serum New Zealand origin (16210064), mouse monoclonal CD81 antibody (10630D), mouse monoclonal CD63 Alexa Fluor^TM^ 488 antibody (MA5-18149), goat-anti-mouse IgG Superclonal^TM^ Alexa Fluor^TM^ 647 antibody (A28181), goat-anti-mouse IgG Superclonal^TM^ Alexa Fluor^TM^ 488 antibody (A28175), goat-anti-mouse IgG Superclonal^TM^ Alexa Fluor^TM^ 555 antibody (A28180), Bovine serum albumin (BSA, BP1600-100), Vybrant^TM^ DiD Cell Labeling Solution (V22887), TetraSpeck microspheres (T72080), and Phalloidin Alexa Fluor™ 647 (A22287) were purchased from Thermo Fisher Scientific. Mouse monoclonal TCR α/β Alexa Fluor™ 647 (306714) was purchased from BioLegend. Ethanol (2701G) was purchased from Decon Laboratories Inc. Paraformaldehyde (15710) and glutaraldehyde (16120) were purchased from Electron Microscopy Sciences. Unconjugated gold colloids (15711–20) were purchased from Ted Pella Inc. ImmunoCult^TM^ Human CD3/CD28 T Cell Activator (10971) was purchased from STEMCELL Technologies.

### Buffers

The following buffers were used for this study. Fixation Buffer: 4% PFA and 0.1% GA in 1x PBS. Post-Fixation Buffer: 4% PFA in 1x PBS. Saponin Wash Buffer: 0.5 % normal goat serum (NGS) and 0.005% saponin in 1x PBS. *d*STORM Buffer: 9.8% (w/v) glucose, 34 µg mL^-1^ catalase, 556 µg mL^-1^ glucose oxidase, 143 mM 2-mercaptoethanol, 50 mM Tris, and 10 mM sodium chloride.

### Cell Culture

Jurkat E6-1 cells (ATCC TIB-152) were cultured in RPMI with 10% Fetal Bovine Serum (FBS) and 100 units/mL penicillin-streptomycin. The cells were kept in a humidified incubator at 37°C supplemented with 5% CO_2_ and split at confluency.

### Sample Preparation

#### Jurkat Activation

200,000 cells were added into a well of a 24-well plate. 5 μL of ImmunoCult^TM^ Human CD3/CD28 T Cell Activator was added to the solution in 200 μL of media containing the cells. The cells were left to incubate for 24 hours in a humidified incubator at 37°C supplemented with 5% CO_2_.

#### Cell Landing

Chambered coverglass wells were coated with 0.01 % Poly-L-Lysine. 50,000 cells were counted and resuspended in PBS. Cells were added to the chambered coverglass wells and incubated for 30 min at 37°C and 5% CO_2_.

#### Fixation and Blocking

The landed cells were fixed in 200 µL of fixation buffer warmed to 37°C for 15 min. The cells were then washed three times with PBS. The cells were blocked and 10% NGS for 1 hour at room temperature. For F-actin and CD81 co-staining, cells were blocked and permeabilized with 0.1% saponin in 10% NGS for 1 hour at room temperature.

#### Immunofluorescence Staining

For single-color CD81 imaging, the cells were primary stained with 10 µg mL^-1^ mouse anti-CD81 in 10% NGS overnight in a humid chamber at 4°C. The cells were washed three times with 1x PBS for 5 minutes. The cells were secondary stained with 5 µg mL^-1^ goat anti-mouse Superclonal Alexa Fluor^TM^ 647 in 10% NGS for 2 hours in a covered humid chamber at room temperature. Following the staining, the cells were washed four times with the 1x PBS. The cells were post-fixed in the post-fixation buffer for 5 minutes, then washed three times with 1x PBS.

For dual-color F-actin and CD81 staining, the cells were primary stained with 10 µg mL^-1^ mouse anti-CD81 in 10% NGS with 0.1% saponin overnight in a humid chamber at 4°C. The cells were washed three times with the saponin wash buffer for 5 minutes. The cells were secondary stained with 5 µg mL^-1^ goat anti-mouse Superclonal^TM^ Alexa Fluor^TM^ 488 in 10% NGS with 0.1% saponin for 2 hours in a covered humid chamber at room temperature. Following the staining, the cells were washed four times with the saponin wash buffer. The cells were post-fixed in the post-fixation buffer for 5 minutes, then washed three times with 1x PBS. The cells were then incubated in 167 nM Phalloidin-Alexa Fluor^TM^ 647 in 10% NGS with 0.1% saponin overnight in a humid chamber at 4°C.

For CD81 and CD63 staining, cells were primary stained with 5 µg mL^-1^ mouse anti-CD81 in 10% NGS overnight in a humid chamber at 4°C. The cells were washed three times with 1x PBS for 5 minutes. The cells were secondary stained with 2.5 µg mL^-1^ goat anti-mouse Superclonal Alexa Fluor^TM^ 647 in 10% NGS for 2 hours in a covered humid chamber at room temperature. Following the staining, the cells were washed four times with the 1x PBS. The cells were post-fixed in the post-fixation buffer for 5 minutes, then washed three times with 1x PBS. The cells were then incubated with 1:100 dilution of mouse-anti-CD63-Alexa Fluor^TM^ 488 overnight at 4°C. The cells were washed three times with 1x PBS for 5 minutes.

For CD81, CD63, and TCR staining, cells were primary with 5 µg mL^-1^ mouse anti-CD81 in 10% NGS overnight in a humid chamber at 4°C. The cells were washed three times with 1x PBS for 5 minutes. The cells were secondary stained with 2.5 µg mL^-1^ goat anti-mouse Superclonal Alexa Fluor^TM^ 647 in 10% NGS for 2 hours in a covered humid chamber at room temperature. Following the staining, the cells were washed four times with the 1x PBS. The cells were post-fixed in the post-fixation buffer for 5 minutes, then washed three times with 1x PBS. The cells were then incubated with 1:100 dilution of mouse-anti-CD63-Alexa Fluor^TM^ 488 overnight at 4°C. The cells were washed three times with 1x PBS for 5 minutes. The cells were then incubated with 2 µg mL^-1^ mouse-anti-TCR-Alexa Fluor^TM^ 647 antibody for 4 hours at room temperature. Following the staining, the cells were washed four times with the 1x PBS.

For membrane staining of control and activated cells, manufacturer’s directions of Vybrant^TM^ DiD Cell Labeling Solution were followed. Cells were then landed on a PLL surface and allowed to attach to the surface for 20 minutes prior to imaging.

### Microscopy

#### TIRF

TIRF microscopy imaging was performed on a Nikon Ti-2 Eclipse inverted microscope coupled with NIS-Elements Advanced Research software for acquisition. A Nikon 100x/1.49 oil immersion objective was combined with 1.5x external magnification to get an effective pixel size of 73 nm. Images were captured in the 488 nm (72 W cm^-2^) and 640 nm (20 W cm^-2^) channels with a Prime 95B sCMOS camera (Serial Number: A18B203004). Images were acquired with a 50 ms exposure time and 16-bit data depth at ∼25°C.

#### DeepSIM

DeepSIM images were acquired using a Nikon Ti-2 Eclipse inverted microscope equipped with a CrestOptics DeepSIM imaging system, a Nikon 60x/1.42 oil immersion objective, and a Kinetix sCMOS camera (Serial Number: A23H723002). NIS-Elements Advanced Research software (Version: 5.42.04) was used for acquisition.

For CD81 and F-actin imaging, the bottom 2 µm of the cell was imaged with a step size of 0.2 µm. The 488 nm and 640 nm laser lines were used with camera exposure times of 200 ms and 50 ms, respectively. All acquisitions utilized the standard imaging mode at room temperature.

For the full-cell volumetric imaging of CD81, 10-14 µm of the cell was imaged using a step size of 0.5 µm. The 640 nm laser lines was used with camera exposure time of 50 ms. All acquisitions utilized the standard imaging mode at room temperature.

For all CD81, CD63, and TCR imaging, only the bottom of the cell was imaged in one plane. The 488 nm, 561 nm, and 640 nm laser lines were used with integration times of 200 ms, 90 ms, and 200 ms, respectively. All acquisitions utilized the Deep imaging mode at room temperature.

#### dSTORM

*d*STORM imaging was performed on a Nikon Ti-2 Eclipse inverted microscope coupled with NIS-Elements Advanced Research software (Version 5.21.03) for acquisition. A Nikon 100x/1.49 oil immersion objective was combined with 1.5x external magnification. A Prime 95B sCMOS camera (Serial Number: A18B203004) was set to 16-bit data depth and 2×2 pixel binning to create a pixel size of 147 nm. Exposure time was set to 20 ms. A region of interest (ROI) of 512×512 pixels (256×256 after binning) was selected. Gold colloids were added prior to imaging as fiducial markers for drift correction. *d*STORM buffer was added into the well of the sample and sealed with parafilm. *d*STORM images were acquired using a 640 nm laser with a measured power density of ∼5 kW cm^-2^ after the objective. 405 nm laser was gradually increased to maintain adequate blinking. 70,000 image frames were obtained at room temperature.

### Data Processing

#### Image Reconstructions

DeepSIM images were reconstructed from the raw data using the Nikon Elements Advanced Research (Version: 5.42.04) software along with the DeepSIM software addition. The reconstructed images (.nd2 file format) were saved and opened in Fiji. For F-actin and CD81 imaging, a Z-projection of all the Z-slices was performed to create a 2D representation of the images.

*d*STORM image reconstruction was performed using the Fiji plugin ThunderSTORM (*50*). Drift correction for T cell imaging was performed using gold nanoparticle fiducial markers. Localized single-molecule events with a localization uncertainty greater than 20 nm and sigma (standard deviation of the integrated Gaussian localization function) greater than 200 nm were filtered out from the analysis. The visualizations shown are normalized Gaussian with a pixel size of 20 nm. No nonlinear or partial image adjustments were performed.

#### Colocalization Analysis

Colocalization analysis was performed in Fiji using the Coloc2 plugin. The background was subtracted using an area outside the cell. ROIs were drawn around the areas of interest. The intensity correlation quotients (ICQ) were calculated and stored in an Excel file.

#### Maxima NND analysis

The images of CD81 and CD63 were compiled, and an ROI was drawn to include only the periphery of the cell, mainly including the microvilli. ImageJ was used to find the maxima in the images of both channels separately. The list was exported in the .csv file format. A custom MATLAB code was employed to calculate the minimum distance between the points. Only non-zero distances were considered for the data sets calculating the CD81 to CD81 and CD63 to CD63 NNDs. All values were compiled by cell for the intracellular comparison, or all together for the histogram visualization.

#### DBSCAN Clustering Analysis

Sections of microvilli from the cells were cropped to isolate x-y coordinate localizations from the *d*STORM reconstruction. The x-y coordinates for each cell’s microvilli were then compiled. DBSCAN analysis was performed using a custom-written MATLAB code that uses the DBSCAN clustering algorithm to cluster the *d*STORM coordinates. The DBSCAN clustering parameters were set to a minimum number of points = 7 and a search radius (Epsilon) = 10. Each cluster identified by DBSCAN had its nearest neighbor distance calculated by comparing the distance of the centroid of the cluster to the centroids of the other clusters in the data set and finding the minimum value. Each cluster’s maximum diameter was calculated by finding the distance between each point in the cluster and finding the maximum value. Area was calculated using boundary function in MATLAB to create a boundary around the cluster and calculating its area. NND, MD, and area values for each cluster were compiled and exported for analysis.

#### Statistical Analysis

Student’s T-tests were performed to assess statistical significance in GraphPad Prism.

## Acknowledgments

The authors thank Hirushi Gunasekara for her valuable comments and assistance in editing the manuscript.

## Author contributions

Conceptualization: YSH, NTR,

HJ Methodology: NTR, HJ, JA

Investigation: NTR

Visualization: NTR

Writing—original draft: NTR

Writing—review & editing: NTR, YSH, HJ, JA

Funding: YSH

## Data availability

All data is available in the main text or supplementary materials. Due to their large file size, raw *d*STORM or SIM images are available upon request. Custom MATLAB scripts are available on GitHub at https://github.com/nrams12/DBSCAN-Clusterer-NND-MaximaCD81.

## Supplementary Materials

Please see Supplementary Materials Document.

