## Supplementary material for "Superresolution Imaging Reveals the Spatial Organization of CD81 Microdomains in Regulating Membrane Signaling on Jurkat T Cell Microvilli": Figs. S1 to S9

Neal T. Ramseier *et al.*

**This PDF file includes:**

Figs. S1 to S9

**A**

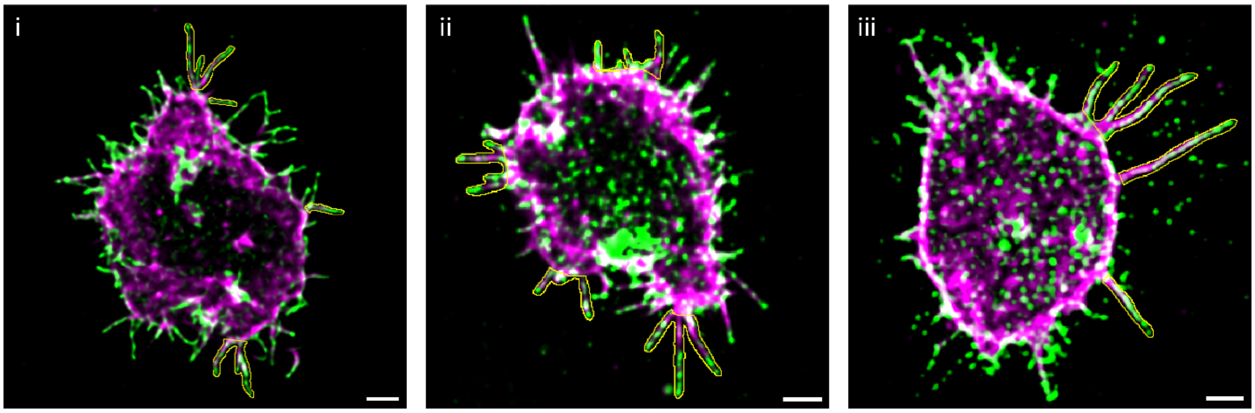


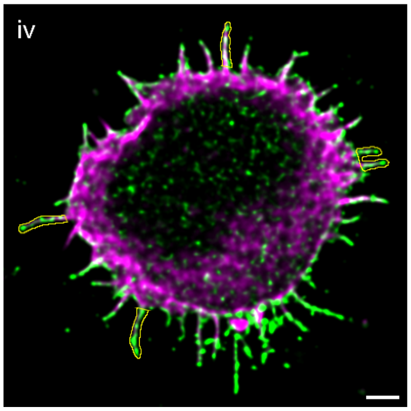


**B**

| Summary Table | ICQ |
|---|---|
| Average | 0.190 |
| Standard Deviation | 0.060 |
| SEM | 0.017 |

**C**

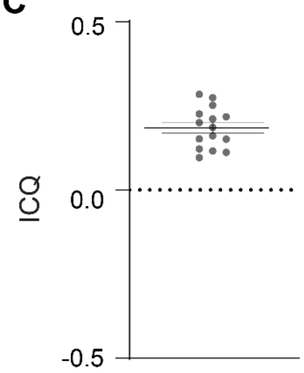


**Figure S1: Quantitative colocalization analysis of CD81 with F-actin.** (**A**) Reconstructed SIM images of CD81 (green) and F-actin (magenta) on a Jurkat T cell. Yellow outlines represent regions of interest utilized for analysis. (**B**) Summary table of the colocalization metrics calculated utilizing the intensity correlation quotients (ICQ) measured along 15 total microvilli from 4 Jurkat T cells. **C**) Distribution of the ICQ values. Bars represent the mean ± standard error of the mean (SEM). Scale bars: 2 μm (**A**).

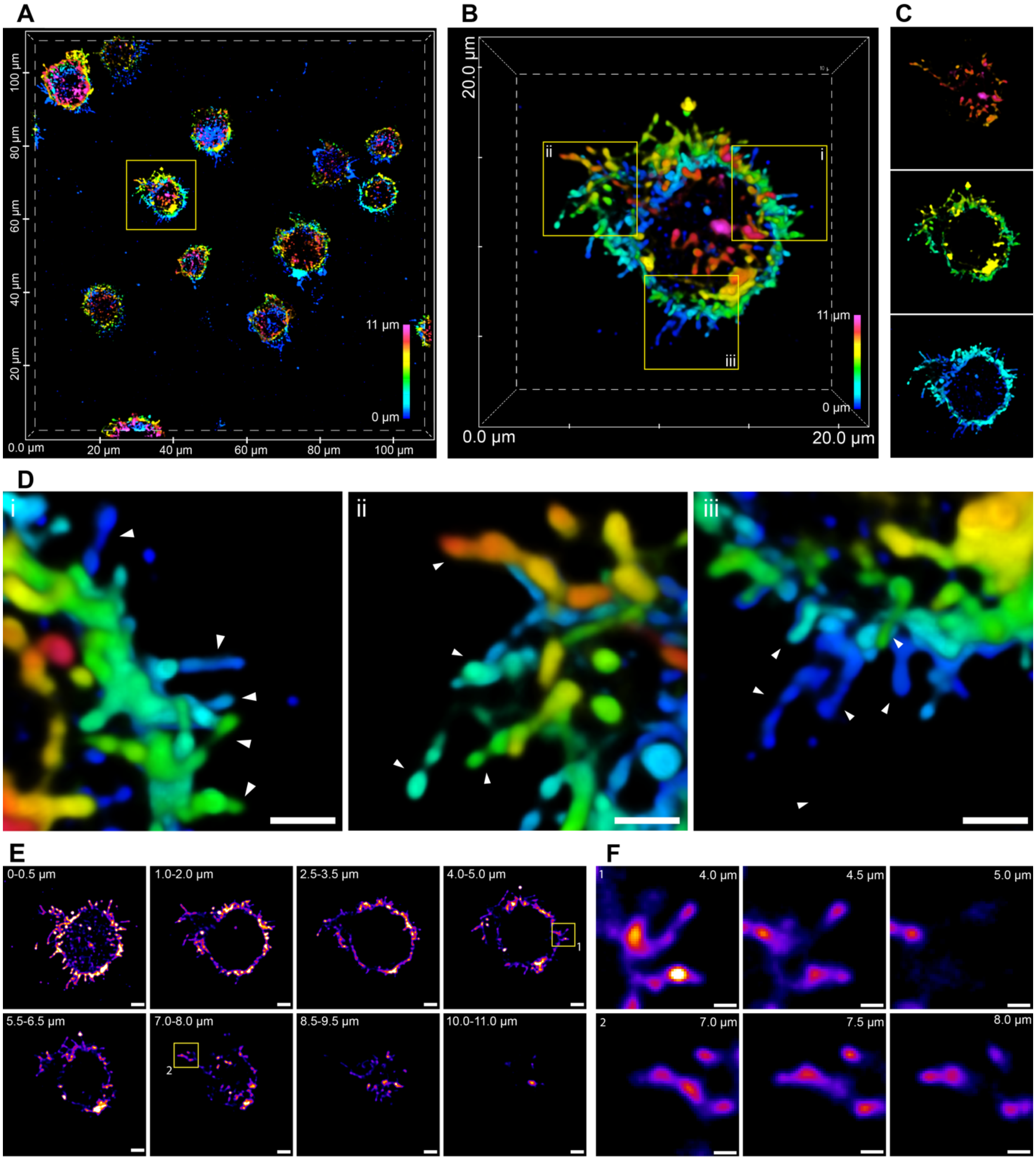


**Figure S2: Volumetric superresolution imaging by DeepSIM revealed the prevalence of CD81 on the microvilli across entire Jurkat T cell surface.** (**A**) Volumetric superresolution image of CD81 across multiple Jurkat T cells shown from the top-down. Color encodes the depth. (**B**) Magnified view of the box region in panel **A**. (**C**) Segmented views of the cell in panel **B** display the volumetric view of the top third (top), middle third (middle), and bottom third (bottom). (**D**) Magnified views of the boxes in panel **B**. Triangles point to locations of microvilli at various *z* positions. (**E**) *z*-slices of the cell in panel **B** displaying the punctated CD81 appearance in the lateral direction at various heights above the coverslip. (**F**) Magnified views of the boxes in panel **E**. Scale bars: 2 µm (**E**), 1 µm (**D**), and 500 nm (**F**).

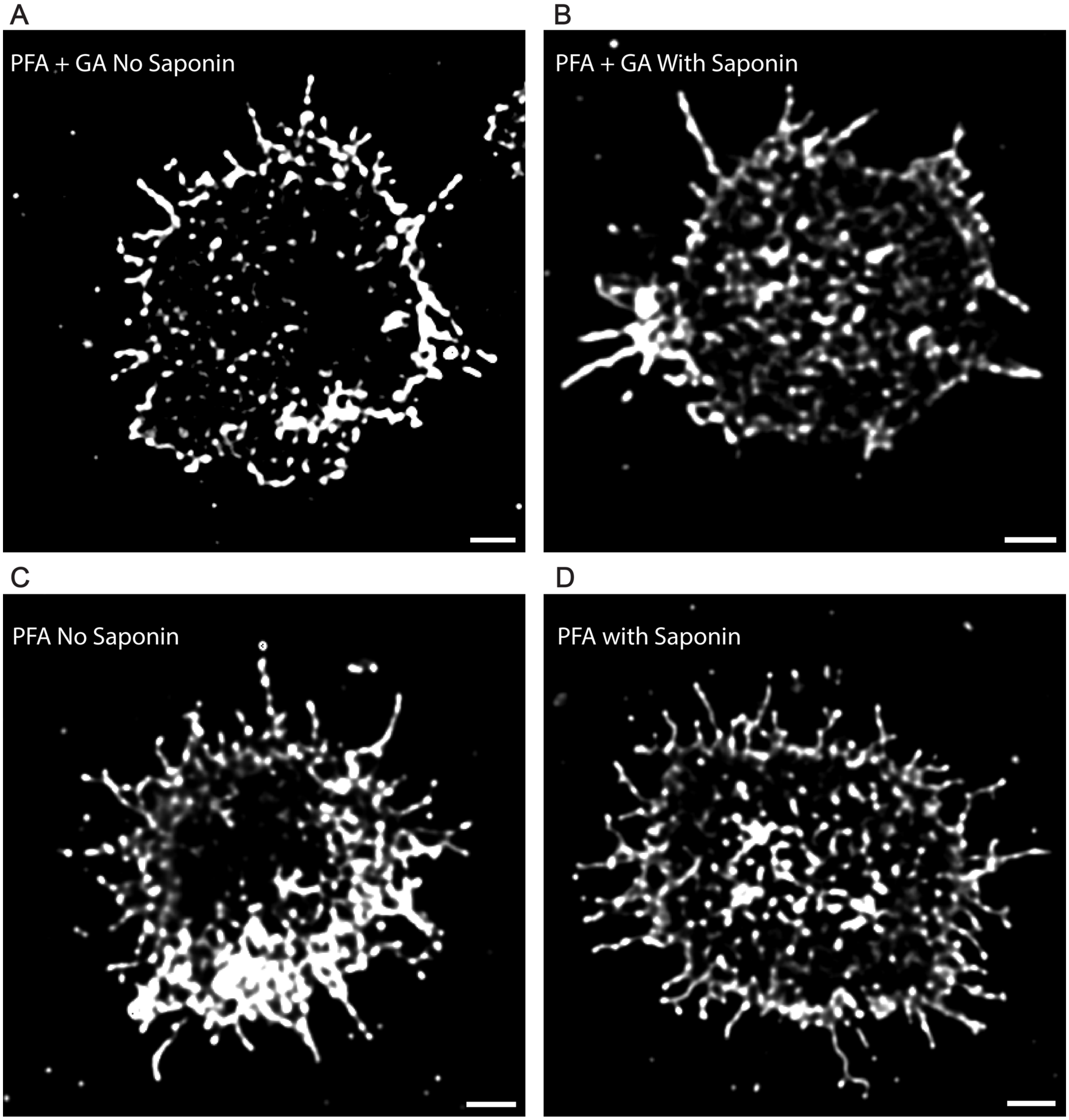


**Figure S3: CD81 intensity puncta observed in various fixation conditions.** (**A**) SIM image of CD81 on a Jurkat T cell fixed with 4% PFA + 0.1% GA without permeabilization. (**B**) SIM image of CD81 on a Jurkat T cell fixed with 4% PFA + 0.1% GA and was permeabilized with 0.1% saponin. (**C**) SIM image of CD81 on a Jurkat T cell fixed with 4% PFA without permeabilization. (**D**) SIM image of CD81 on a Jurkat T cell fixed with 4% PFA and was permeabilized with 0.1% saponin. Scale bars: 2 µm.

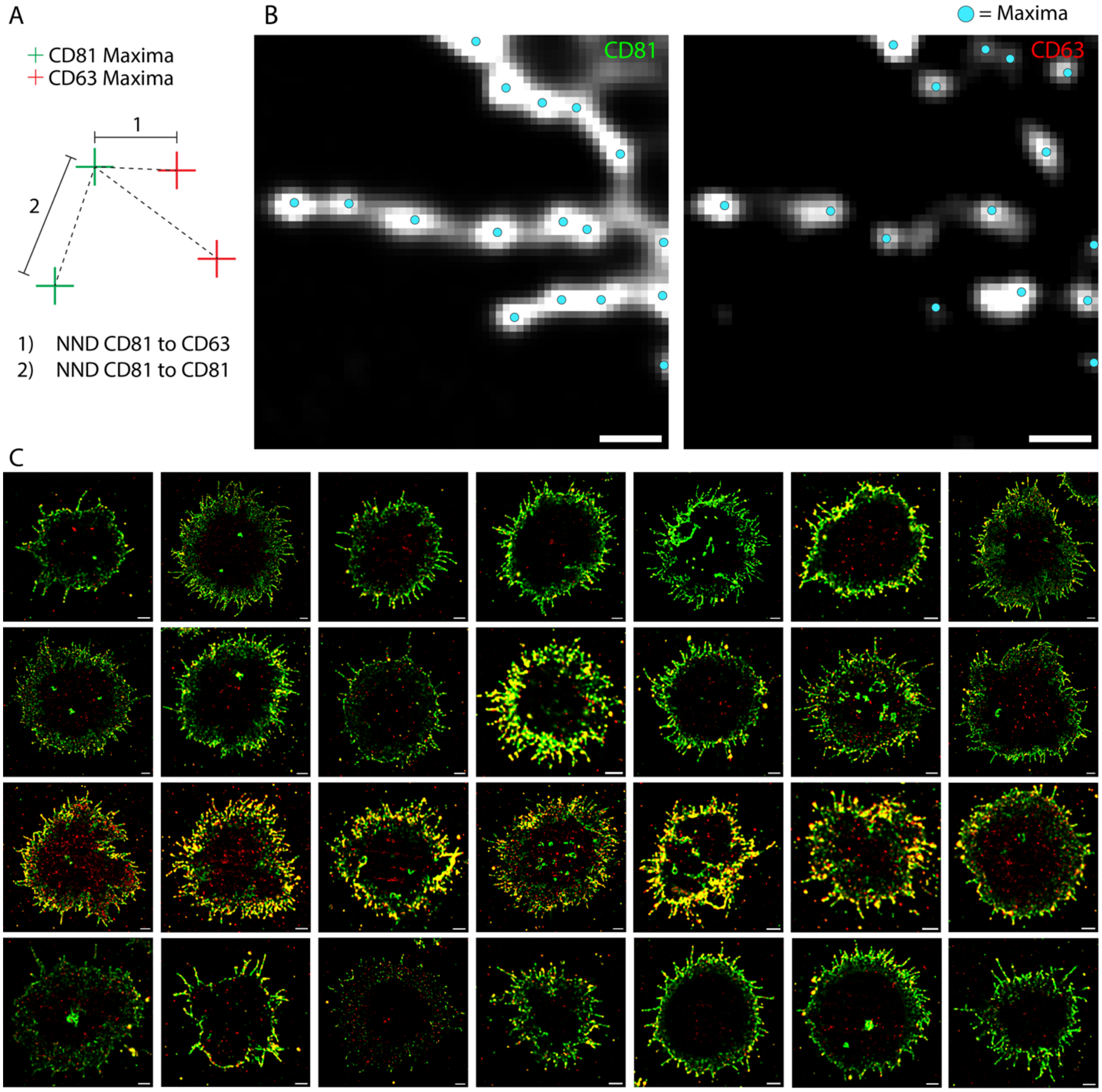


**Figure S4: Nearest neighbor distance (NND) of the pixel intensity maxima analysis.** (**A**) Schematic depicting the NND calculation between CD81 maxima and CD63 maxima. (**B**) Sample region of CD81 (left) and CD63 (right) with maxima shown as cyan circles. (**C**) 28 SIM images of CD81 (green) and CD63 (red) included in the NND maxima analysis. Scale bars: 2 µm (**C**) and 500 nm (**B**).

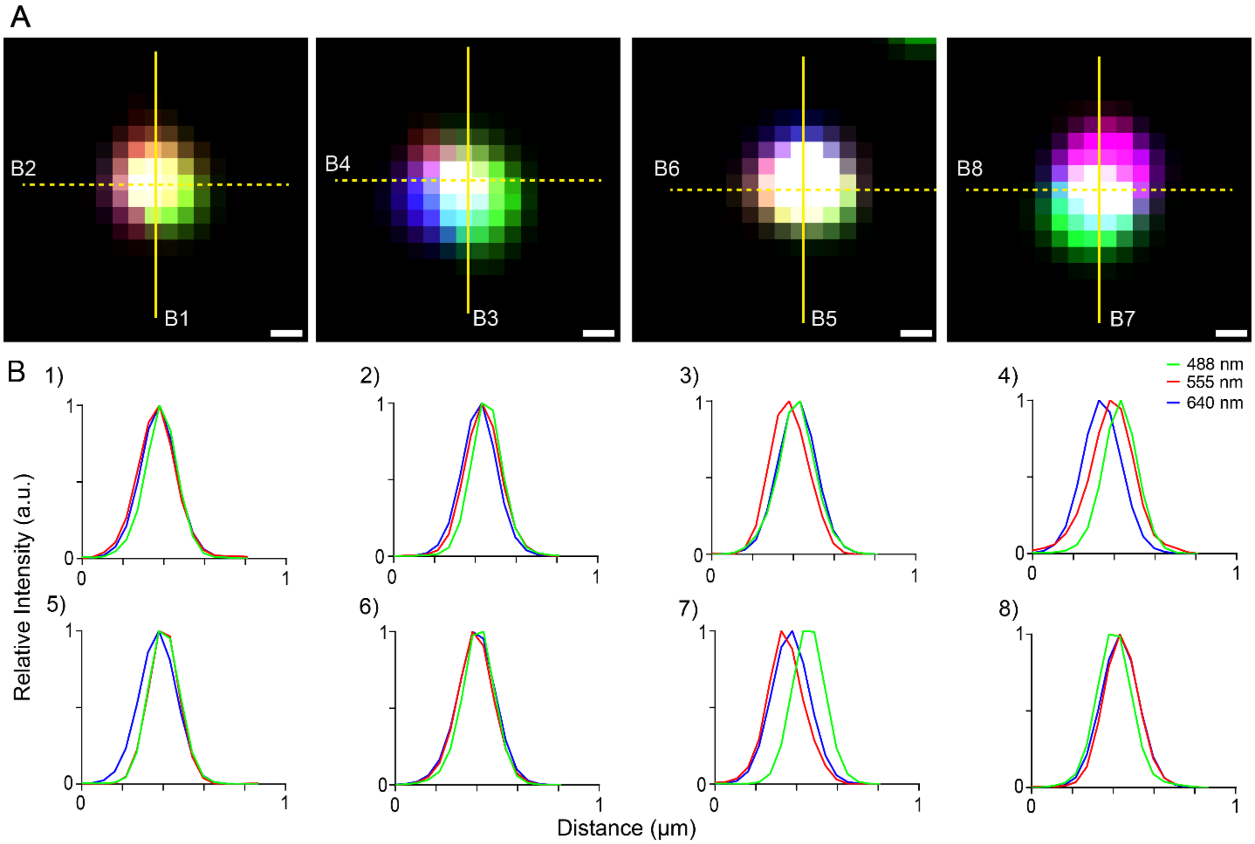


**Figure S5: SIM images of 0.2 µm TetraSpeck™ microspheres reveal varying chromatic shifts between channels.** (**A**) SIM images of TetraSpeck™ microspheres in the 488 nm (green), 561 nm (red), and 640 nm (blue) channels. (**B**) Line profiles displaying the relative intensity of all three channels corresponding to the lines in panel **A.** Scale bars: 100 nm (**A**).

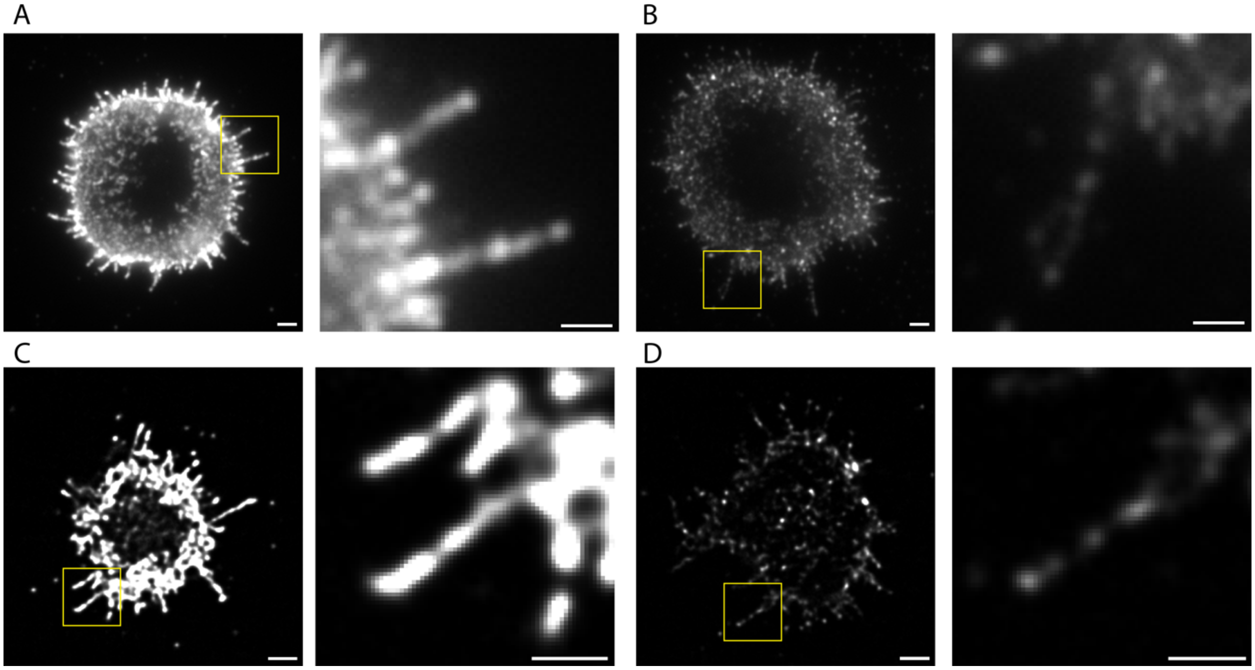


**Figure S6:** TIRF images of CD81 on control (**A**) and 24-h activated (**B**) Jurkat T cells with magnified views of the boxed region shown on the right. Images are displayed with the same contrast. SIM images of CD81 on control (**C**) and 24-h activated (**D**) Jurkat T cells with magnified views of the boxed region shown on the right. Images are displayed with the same contrast. Scale bars: 2 µm (**A-D(left)**) and 1 µm (**A-D(right)**).

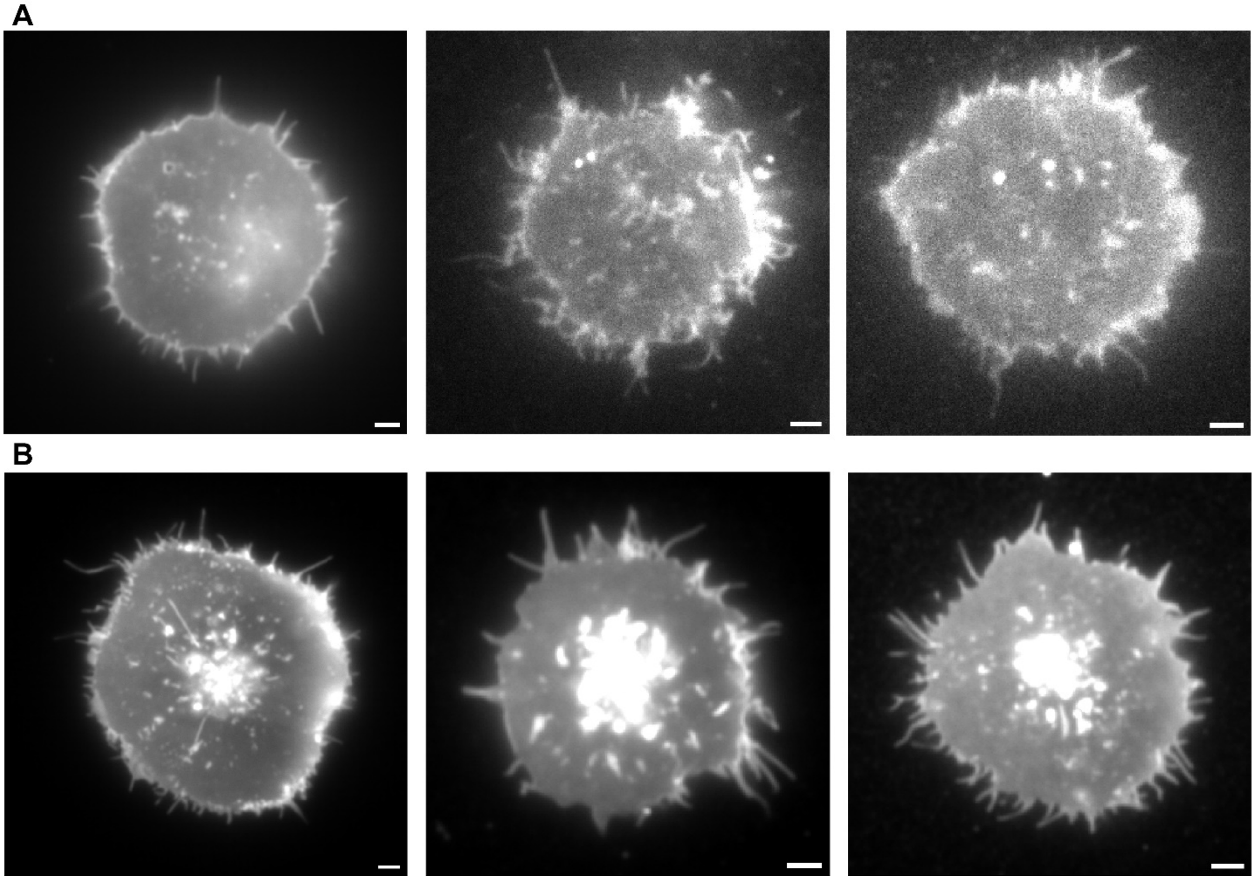


**Figure S7:** (**A**) 24-h activated Jurkat T cells labeled with a membrane dye. (**B**) Control Jurkat T cells labeled with a membrane dye. Scale bars: 2 µm.

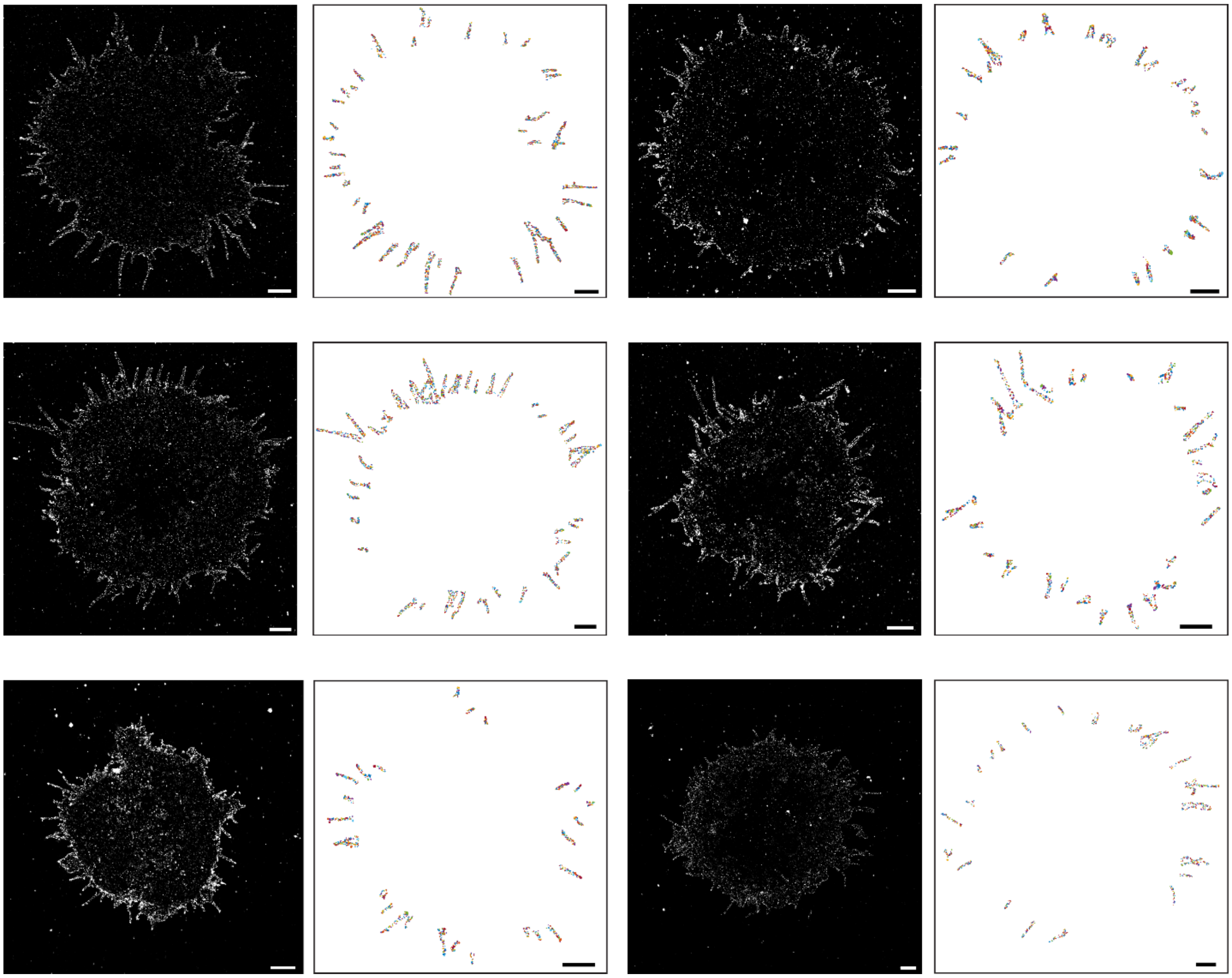

**Figure S8:** *d*STORM images of CD81 across control Jurkat T cells (left) and DBSCAN analysis used to calculate the average cluster nearest-neighbor distance (NND), maximum diameter (MD), and area for each cell (right). Scale bars: 2 µm.

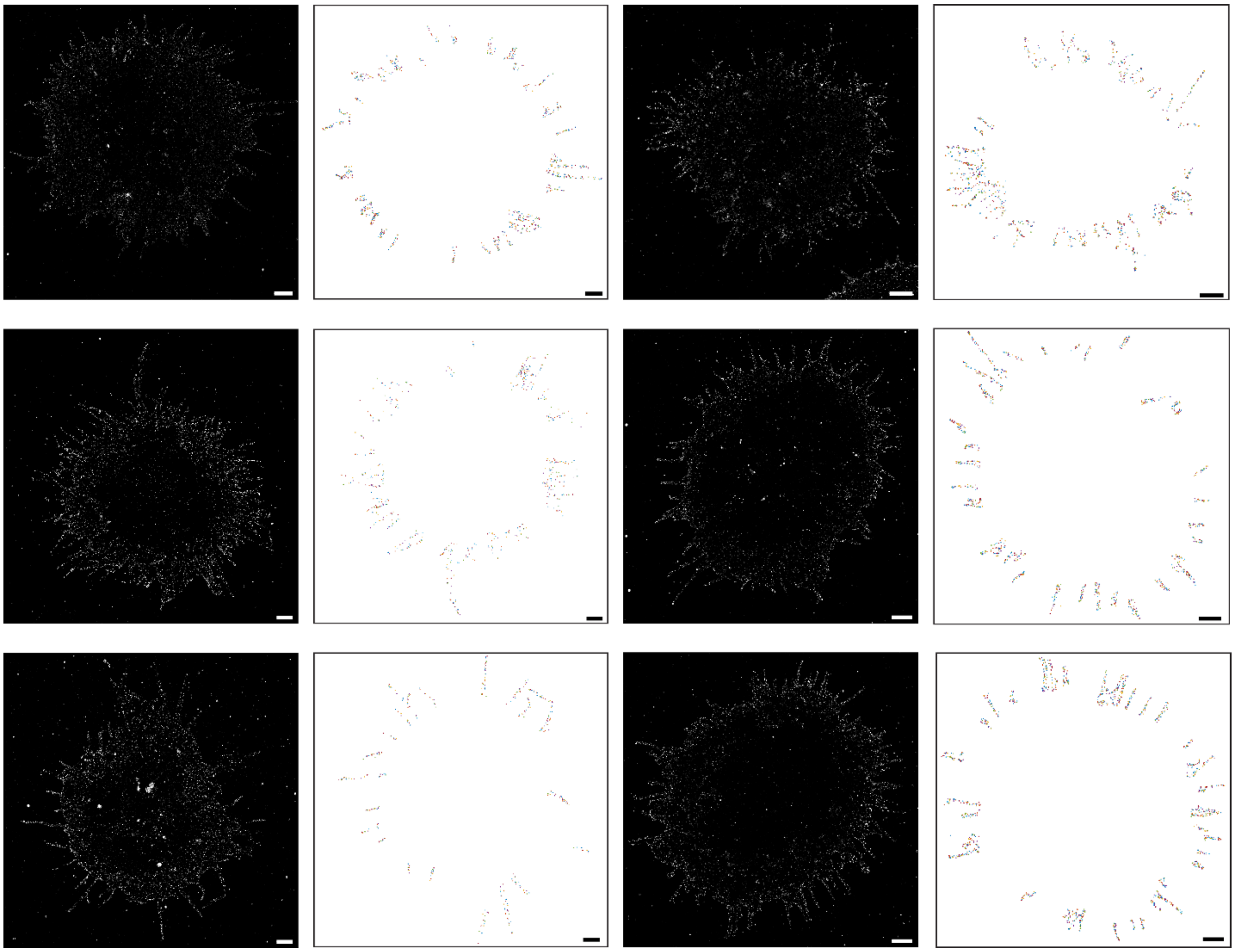

**Figure S9:** *d*STORM images of CD81 across 24-h activated Jurkat T cells (left) and DBSCAN analysis used to calculate the average cluster nearest-neighbor distance (NND), maximum diameter (MD), and area for each cell (right). Scale bars: 2 μm.
